# *Vibrio harveyi* plasmids as drivers of virulence in barramundi (*Lates calcarifer*)?

**DOI:** 10.1101/2025.02.05.636614

**Authors:** Roisin Sullivan, Joy A Becker, Ruth N Zadoks, Carola Venturini, Ana I. S. Esteves, Suresh Benedict, Dani L Fornarino, Hannah Andrews, God’spower R Okoh, Vidya Bhardwaj, Mark Sistrom, Mark E Westman, Nguyen Ngoc Phuoc, Francisca Samsing

## Abstract

*Vibrio* species are an emerging public and animal health risk in marine environments and the opportunistic bacterial pathogen *Vibrio harveyi* is a major disease risk for tropical aquaculture. Current understanding of virulence in *V. harveyi* is limited by strain-specific variability and complex host-pathogen dynamics. This study sought to integrate genomic investigation, phenotypic characterisation and *in vivo* challenge trials in barramundi (*Lates calcarifer)* to increase our understanding of *V. harveyi* virulence. We identified two hypervirulent isolates, Vh-14 and Vh-15 that caused 100% mortality in fish within 48 hours, and that were phenotypically and genotypically distinct from other *V. harveyi* isolates. Virulent isolates contained multiple plasmids, including a 105,412 bp conjugative plasmid with type III secretion system genes originally identified in *Yersinia pestis*. The emergence of this hypervirulent plasmid-mediated patho-variant poses a potential threat to the sustainable production of marine finfish in Southeast Asia, the Mediterranean and Australia. In addition, we observed an effect of temperature on phenotypic indicators of virulence with an increase in activity at 28°C and 34°C compared to 22°C. This suggests that temperature fluctuations associated with climate change may act as a stressor on bacteria, increasing virulence gene secretion and host adaptation. Our results utilising a myriad of technologies and tools, highlights the importance of a holistic view to virulence characterisation.

## 1. Introduction

The genus *Vibrio* encompasses a broad group of opportunistic and highly pathogenic marine bacteria which are anticipated to increase in prevalence and distribution as climate change continues [1]. There are more than 70 described *Vibrio* spp. which are considered ubiquitous in marine and brackish water worldwide and can cause disease in a wide array of species including humans, finfish, crustaceans and molluscs [2]. Whilst *Vibrio* spp. can exist in a wide-range of thermal conditions, they typically thrive in warmer environments and as average sea surface temperatures are increasing worldwide, so too has the extent and prevalence of *Vibrio* spp. globally [3]. In the past decade, there has been a sustained emergence and detection of *Vibrio* spp. in previously unaffected regions and outside expected detection windows directly in relation to increasing sea surface temperatures and changing climatic conditions [4]. Notable examples include the 2014 incursion of non-O1 *V. cholera* into subarctic regions of Sweden and Finland [5], the first detection of non-O1 *V. cholera* in British Columbia, Canada, following marine heat-waves in 2018 [3], and the increase in *V. vulnificus* infections associated with Hurricane Ian’s landfall in Florida in 2022 [6].

Whilst there is a clear and direct public health risk for humans with increased incursions and detections of pathogenic *Vibrio* spp., they also pose a major barrier to the expansion of aquaculture and the health of farmed and wild aquatic organisms [7]. Similar to the incursion of human pathogenic non-O1 *V. cholera* into previously unaffected regions, the extent and occurrence of *Vibrio* spp. that are pathogenic for aquaculture species is also increasing. Alongside new incursions and the prolonged detection of *V. rotiferianus* and *V. jasicida* in shellfish in Wales and England [8], increasing reports of *V. harveyi* are occurring in regions of the Mediterranean, with *V. harveyi* now considered one of the major disease threats to European seabass (*Dicentrarchus labrax*) and gilthead seabream (*Sparus aurata*) production [9, 10]. This trend is echoed in Southeast Asia, with production losses due to vibriosis accounting for more than 7% of production costs in sea-cage barramundi in Malaysia [11]. In China, which accounts for over 30% of global aquatic animal production [12], *V. harveyi* was the dominant bacterial pathogen identified in diseased marine finfish farmed in the southern Guangdong and Hainan regions in 2020 [13]. This echoed earlier findings which identified *V. harveyi* as the dominant pathogen in diseased finfish sampled from 2015 to 2018 in China, Malaysia and Vietnam [14].

Whilst *V. harveyi* is typically more opportunistic than other *Vibrio* spp., such as *V. vulnificus* or *V. alginolyticus,* it has increasingly been detected as the causative agent of disease in a wide range of host species including seahorses (*Hippocampus kuda*) [15], European abalone (*Haliotis tuberculate*) [16], mud crabs (*Scylla* spp.) [17], prawns [18] and numerous finfish species [19]. Clinical signs of disease can be highly varied and species-specific. In finfish, vibriosis associated with *V. harveyi* commonly manifests as gastroenteritis, muscle necrosis, fin erosion and skin lesions [20]. This heterogeneity in clinical manifestations of infection may in part be due to the fact that *V. harveyi* may be reliant on co-stressors or host compromise for disease to occur [21]. To date, the study of virulence in *V. harveyi* has mainly focused on PCR-based analysis of known and atypical virulence genes from other *Vibrio* spp. such as *flaC* from *V. anguillarum* and *ctxA* from *V. cholerae* [13, 22]. Although several whole genome investigations have been performed for *V. harveyi* [10, 23, 24], a holistic understanding of virulence in *V. harveyi* is still lacking – particularly in comparison to *V. cholerae* [25, 26], and *V. parahaemolyticus* [27].

This study aims to advance our understanding of virulence in *V. harveyi* by combining *in silico* genomic characterisation with traditional *in vitro* phenotypic analysis and *in vivo* challenges using juvenile barramundi (*Lates calcarifer*) as a model tropical fish.

## 2. Methods

### 2.1 Sample collection and bacterial isolation

Isolates from diseased fish were obtained from Australia in 2022 and Vietnam in 2023. Australian *Vibrio* isolates originated from clinically moribund farmed barramundi (*Lates calcarifer)* that were submitted to Berrimah Veterinary Laboratory, Northern Territory (BVL) for culture and identification of bacterial pathogens. Upon arrival at BVL, an eye and three skin swabs post handling and/or grading (n = 4) were inoculated onto marine agar (Difco 2216), tryptone soy agar with 5% sheep’s blood (ThermoFisher Scientific, Australia PP2166) and thiosulfate-citrate-bile salts-sucrose (TCBS) agar (ThermoFisher PP2013) and incubated at 26°C until colony growth was observed. Vietnamese isolates were collected in June 2023, during a syndromic surveillance study in marine polyculture seacage farms in the Quang Ning Province of Ha Long Bay (Vietnam). Using selective media (TCBS agar plates), Vibrio *spp.* were isolated from moribund fish showing gross external pathology consistent with bacterial infections (scale loss and skin ulcers, and reddening at the base of the fins and ventral areas of the body). Fish were humanely euthanised with a rapid blow to the head and sampled by aseptically swabbing the head-kidney with a 1 μL inoculating loop. Eight *Vibrio* spp., displaying yellow colonies on TCBS plates, were identified as *Vibrio harveyi* by matrix-assisted laser desorption-ionization time of flight mass spectrometry (MALDI-ToF MS; ThermoFisher Scientific). These isolates came from six out of thirty fish sampled during this study and included spotted scat (*Scatophagus argus*) (n = 1), rabbitfish (*Siganus gutatus*) (n = 2) and grouper (*Epinephelus fuscoguttatus*) (n = 3). The eight isolates were passaged onto marine agar (Difco 2216) twice and incubated at 28°C to obtain pure colonies. All twelve suspected *V. harveyi* isolates (n = 4 collected from Australia and n = 8 collected from Vietnam) were identified to species level by MALDI-Tof MS [28] before sequencing as described in Section 2.2.

### 2.2 Virulence typing *in silico*

Bacterial DNA was extracted using the MagMAX CORE Nucleic Acid Purification Kit (327-000) on the KingFisher Flex System (ThermoFisher Scientific) or with the Quick-DNA Miniprep Plus Kit (ZymoResearch D4068, USA) according to the manufacturer’s protocol for the isolates, respectively. DNA quality was checked with a Qubit^TM^ fluorometer (ThermoFisher Scientific). Following extraction, short-read libraries were prepared with the Illumina DNA Prep kit and sequenced as 300-bp paired-end reads on an Illumina MiSeq^TM^ (Illumina, USA) at BVL to a depth of at least 60x coverage. Following sequencing, genomes were *de novo* assembled using SPAdes v3.15.5 [29] with default parameters and annotated with Prokka v1.14.1 [30]. After assembly, the average nucleotide identity (ANI) of the genomes was assessed using FastANI v1.32 [31] against the NCBI reference genomes for *V. harveyi* (SB1, NCBI: PRJNA972608) and *V. campbelli* (BoB- 53, NCBI: PRJNA429202) because of high phenotypic and genomic similarities between the species [32, 33].

Long-read sequencing was performed for the Vietnamese isolates with a R10.4.1 flow cell (FLO- MIN114) on a MinION^TM^ Mk1C (Oxford Nanopore Technologies, UK) and libraries were prepared with the Rapid Barcoding Kit V14 (SQK-RBK114). Following sequencing, POD5 files were basecalled in MinKNOW v.24.02.8 using Dorado super accuracy basecaller model (SUP; dna_r10.4.1_e8.2_400bps_5khz_sup) enabling barcode and adapter trimming at basecalling. Raw fastq files were then assembled using the Genomics in a Backpack pipeline [34] developed by the Sydney Informatics Hub in collaboration with the Sydney School of Veterinary Science (https://github.com/Sydney-Informatics-Hub/ONT-bacpac-nf). Briefly, the quality of raw reads was assessed using NanoPlot v.1.42.0 [35] and PycoQC v2.5.2 [36]. Reads were then screened with Kraken2 v2.1.3 [37] to check for contamination prior to assembly. Where possible a hybrid assembly was performed by generating a consensus assembly using Flye v2.9.3 [38, 39] and Unicycler v0.4.8 [40]. If the two long-read assemblies could not be hybridised, only the Flye assembly was kept. For isolates which contained plasmids, the plasmids were assembled using Plassembler v1.6.2 [41]. Following consensus assembly, all contigs were annotated with Bakta v1.9.2 [42]. Antimicrobial resistance (AMR) genes in the long-read assemblies were found using AMRFinderPlus v3.12.8 [43] and the presence of virulence genes was assessed using Abricate (https://github.com/tseemann/abricate) and the virulence factor database (VFDB) (doi:10.1093/nar/gkv1239). Prophage content of the genomes was predicted in the long-read assemblies using PHASTEST v3.0 [44].

To investigate virulence *in silico*, the protein-coding sequences of virulence genes commonly described in *V. harveyi*, or identified in other *Vibrio* species (atypical to *V. harveyi*) were curated into a localised BLAST database for BLASTx analysis of the Prokka-annotated short-read assemblies. In this analysis, publicly available *V. harveyi* genomes were also included (see below under phylogenetic investigation). Forty-one virulence genes were included as identified in prior studies [22, 45–49] and an additional eight genes were included following manual identification *in silico* and via Abricate VFDB output resulting in 49 protein-coding sequences for investigation (S1 Table). All confirmed *V. harveyi* genomes included for phylogenetic analysis, including the NCBI reference genome (SB1), were analysed using blast+/2.12.0 with flag-task blastx and - evalue 0.00001. Furthermore, for the plasmids identified via long-read sequencing using Plassembler in Vh-14 and Vh-15, an additional BLASTx was performed as described above against the protein-coding sequences for the reference genomes of *Yersinia enterocolitica* (Y11, NCBI: SAMEA2271938)*, Y. pestis* (A1122, NCBI: SAMN02603531) and *Y. pseudotuberculosis* (NCTC10275, NCBI: SAMEA4442458) following the identification of *ypkA* in plasmids of these isolates. Visualisation of the plasmids was performed using Proksee [50] and the gggenes package in R [51] for further visualisation of the locus of interest.

Phylogenetic investigation of the isolates was conducted using Snippy v4.1.0 [52] by identifying single nucleotide polymorphisms (SNPs) in each isolate against the *V. harveyi* reference genome (SB1) and generating a core genome SNP alignment with command snippy-core. For comparative analysis, publicly available genomes were identified and screened in the National Centre for Biotechnology Information (NCBI) Sequence Read Archive (S2 Table). Genomes were included if isolates originated from vertebrates or environmental samples, included geographical and isolation source data, were sequenced on an Illumina platform and fastq files were readily available. Genomes (n = 28) were downloaded and annotated using Prokka as above. Following SNP identification for all genomes, a recombination free-alignment was created using Gubbins v3.3.5 [53] and a maximum-likelihood phylogeny constructed in IQ-TREE v1.6.12 [54] using a GTR model and 1000 bootstrap replicates [55]. The tree was then annotated in FigTree v1.4.4 [56] and Microreact [57].

### 2.3 Virulence typing *in vitro*

For the *V. harveyi* isolates collected from moribund fish in Vietnam and *V. harveyi* isolate TCFB- 0558 (NCBI: PRJNA940306) [21], eight phenotypic assays were performed as well as infrared biotyping using Fourier Transform infrared spectroscopy (Fig 1). All isolates were grown aerobically from cryo-preserved frozen aliquots on marine-agar (Difco 2216) at 28°C overnight, and passaged onto a new marine agar plate and incubated aerobically overnight at 28°C. This subculture was then used for testing of lipase, phospholipase, caseinase, haemolysis and urease activity. For each assay, each isolate was stabbed into three different plates to ensure three replicates, and this was repeated for three temperatures: 22°C, 28°C and 34°C. For each plate-based assay, *E. coli* (ATCC 11775) was used as a negative control. For urease, *Yersinia enterocolitica* (ATCC 23715) was used as a positive control and for the other tests, *P. aeruginosa* (NCTC 10662) was used as the positive control. For the other phenotypic assays, one colony of each subculture was inoculated into marine broth (Difco 2216) and incubated overnight at 28°C. Density of the overnight broth cultures was standardized by diluting the suspensions in sterile PBS to an OD_600_ of 0.5 before testing swarming, gelatinase and biofilm formation activity. For gelatinase and biofilm testing, each isolate was inoculated into three separate tubes or plates to ensure three replicates, and this was repeated for each temperature. To evaluate swarming, five replicates were performed for each isolate at each temperature. For gelatinase, *E. coli* (ATCC 11775) was used as a negative control and for biofilm formation, uninoculated marine broth was used as the negative control. *P. aeruginosa* (NCTC 10662) was used as the positive control for each tube-based test. A detailed description of each test, including composition of the media, is provided in S1 Appendix. Plates were observed every 24 hours and activity zones were measured using a ruler at 24, 48 or 72 hours as specified in S1 Appendix. This was true for all assays with the exception of urease (read at 48 hours only), biofilm formation (peg-based assay, see S1 Appendix) and infrared biotyping. For infrared biotyping, isolates were grown on marine agar from frozen aliquots and incubated for 24 hours at 28°C, isolates were then sub-cultured on marine agar and incubated again for 24 hours at 28°C. From this second passage, colonies were collected for biotyping and to ensure sufficient biological replicates, sub-culture from the original passage was repeated and incubated for 24 hours at 28°C.

**Fig 1.**
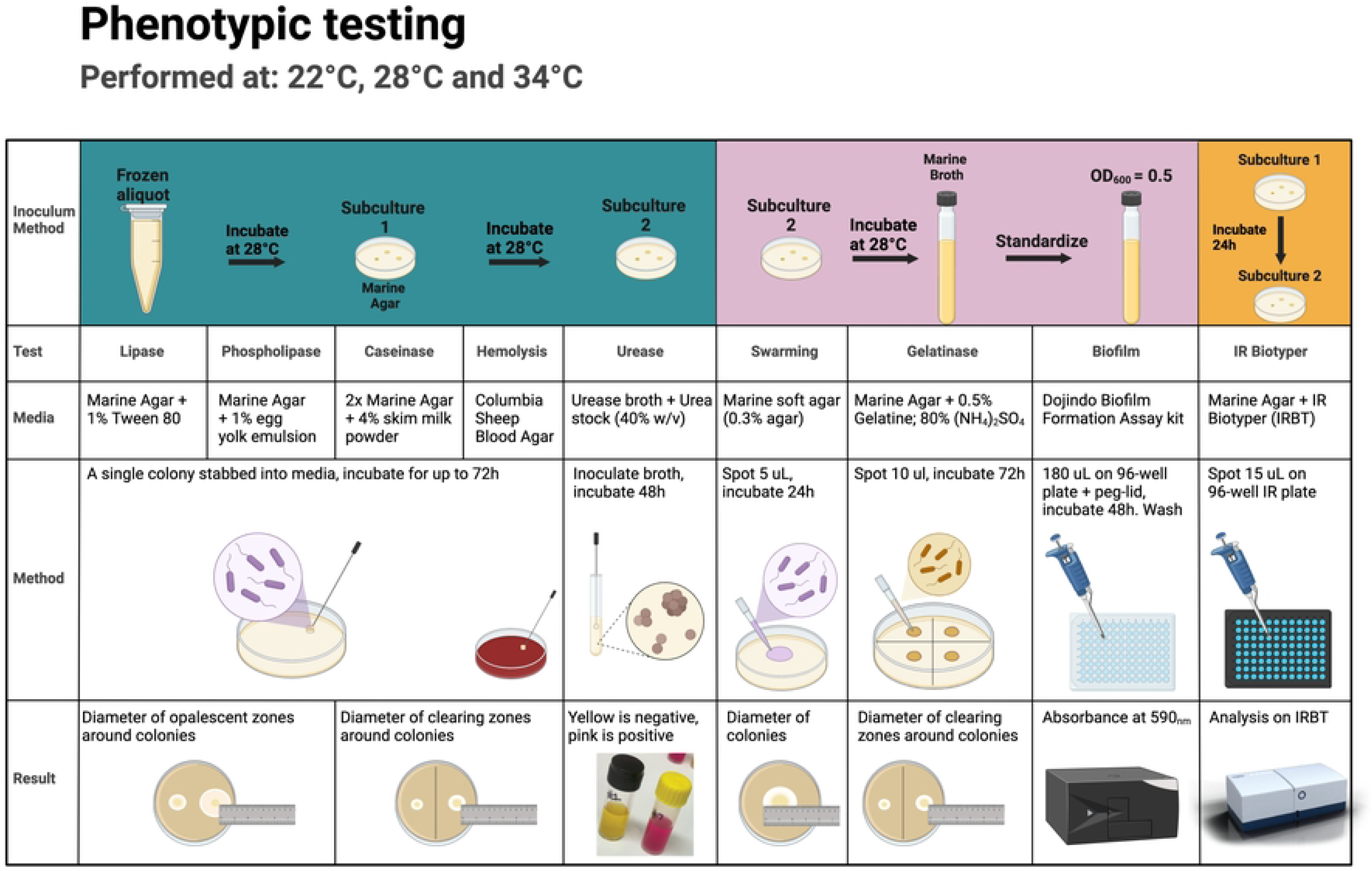
Graphical summary of phenotypic tests performed for *Vibrio harveyi* isolates. Graphical summary of phenotypic tests (lipase, phospholipase, caseinase, haemolysis, urease, swarming, gelatinase, biofilm formation and infrared (IR) biotyping performed for *Vibrio harveyi* isolates Vh-14, Vh-15, Vh-26, Vh-28, Vh-31, Vh-33, Vh-47 and TCFB-0558 and positive and negative assay controls. Swarming was tested using five replicates, all other assays were performed in triplicate, and conducted at 22°C, 28°C or 34°C. Created in BioRender. Sullivan, R. (2025) https://BioRender.com/k48o607.

### 2.4 Pathogenicity *in vivo*

#### 2.4.1 Fish care and maintenance

Experimental challenge studies were performed in juvenile barramundi to complement *in silico* and *in vitro* virulence characterisation. Fish were purchased from a commercial hatchery (Mainstream Aquaculture, Werribee, Victoria) and acclimated for at least two weeks prior to experimental challenges. Three hundred and fifty fish (∼ 0.5 g) were divided into three 500 L circular tanks containing 350 L of seawater (30 ppt) (Red Sea Salt, Red Sea Aquatics) and fed a commercial diet (Otohime C2, BioKyowa, USA) at 8% body weight twice daily for two weeks during the acclimation period. Grading was performed weekly, and fish were grouped based on size to reduce cannibalism. Fish were kept under a 12 hour light:dark schedule and water quality was monitored daily with temperature maintained at 28 ± 1°C, pH at 8.0 ± 0.2 and dissolved oxygen above 6 mg L^−1^. Water quality parameters were monitored using colorimetric test kits (API Saltwater Master Test Kit) and partial water changes were performed as required to maintain total ammonia (<1.0 mg L^−1^), nitrite (<0.5 mg L^−1^) and nitrate (<80 mg L^−1^) at specified levels.

An experimental trial (Experiment 1) was conducted to assess virulence of the *V. harveyi* isolates *in vivo* with two biological replicates from the same cohort of fish. In Replicate 1, fish had a mean weight of 4.5 g ± 1.7 g (mean + SD) and a total length of 74.4 mm ± 8.5 mm, and in Replicate 2, fish had a mean weight of 11.9 g ± 3.4 g and a total length of 103.8 ± 10.0 mm. All fish used in the experimental challenge trials were apparently healthy based on visual assessment of external and internal gross pathology and the absence of systemic bacterial infection in the population was confirmed by random sampling of three fish from each of the holding tanks and plating of a head-kidney swab onto TCBS and tryptone soy agar with 5% sheep blood (ThermoFisher PP2166). Plates were incubated for up to 72 hours at 28°C to confirm the absence of bacterial colonies.

Following confirmation of the health status of fish in the holding tanks, fish were hand netted and transferred to the experimental aquaria for five days of acclimation prior to challenge. Experimental aquaria (100 L) contained 90 L of seawater and water quality was monitored daily to maintain conditions. Each experimental tank was set-up with a biofilter and filtration unit (Aquaclear Power 285 Filter 110), an airstone and air pump (Aqua One stellar 200D Air Pump) for aeration, a heater (Aqua One 200W quartz heater) to maintain water temperature, and a UV steriliser (AA Aquarium, 12V GKM24W). Water quality was monitored daily and maintained as above, the room was temperature-controlled with an air-conditioning system (Hyper Inverter, Mitsubishi Heavy Industries, Australia) to assist in maintaining water temperature. Each tank contained 10 fish to minimise inter-fish aggression.

#### 2.4.2 Challenge trials

Experiment 1 (Replicates 1 and 2) involved randomly assigning nine tanks to one of seven *V. harveyi* isolates collected by the University of Sydney, a known virulent *V. harveyi* isolate (TCFB- 0558; [21]), or a negative control (sham-inoculated with PBS). Replication (n = 2) for each isolate was achieved by replicating the experiment over time rather than using spatial replication. Using a challenge concentration of approximately 3.5 × 10^8^ CFU mL^-1^, based on previous findings for isolate TCFB-0558 [21], we assessed whether these isolates could cause mortality in in juvenile barramundi without additional stressors. For the isolate with the highest and most rapid mortality across the two replicates, a dose-titration experiment was subsequently conducted (Experiment 2). Inoculum preparation was performed according to Samsing [21]. Briefly, isolates were grown as a lawn on marine agar at 28°C. The suspensions were harvested, resuspended, homogenized and diluted to an OD_600_ of ca. 0.250 that was estimated to be 3.5 × 10^8^ CFU mL^-1^. The CFU count was retrospectively determined using the drop count method [58] on 3% marine agar to minimise swarming. Dilutions were only counted if they contained between 5-80 colonies per drop to minimize variability.

For Experiment 1, fish were anaesthetised with benzocaine (60 mg L^-1^) and injected intra-muscularly (IM) in the left flank directly under the dorsal fin with 50 μL of adjusted *V. harveyi* inoculum in PBS giving a final inoculum per fish ranging from 1.4 - 2.2 × 10^7^ CFU fish^-1^, or 50 μL of sterile PBS (S3 Table). Fish were monitored twice a day for seven days post infection. Any fish showing severe clinical disease (e.g. lethargy, weak or erratic swimming, keeling on the side and minimal response to stimulus) were euthanised with an overdose of benzocaine (>150 mg L^- 1^), and counted as a mortality for analysis in accordance with animal welfare and ethics requirements of the project. During the experimental period, all moribund and dead fish were sampled to confirm *V. harveyi* infection and the head-kidney swabbed with a 1 μL inoculating loop onto TCBS plates. For fish with visible skin lesions, the caudal margins of the lesion were also swabbed and streaked onto plates. On Day 8, the remaining fish in each tank were euthanised, and up to 4 fish per tank (depending on number of fish left in each tank) were sampled at random to assess infection by culturing head-kidney swabs on TCBS. All TCBS plates were incubated for up to 48 hours at 28°C and inspected for growth. To confirm *V. harveyi* infection, colony PCR was performed on a pool of 4-6 colonies from each plate. Colonies were collected using a 10 μL loop and resuspended in 300 μL of sterile PBS and flick mixed before being heated at 56°C for 10 minutes to lyse the cells. This was then centrifuged at 17,000 *g* for 10 minutes and the supernatant transferred to a new tube and stored at −20°C until required. This supernatant was used as the template for a conventional PCR using the *toxR* gene primers (382 bp product) from Pang [59]. Briefly, a 12.5 μL reaction was prepared containing 1 μL of template DNA, 2.5 μL of 5x MyTaq^TM^ Red Reaction buffer, 0.25 μL of MyTaq Red DNA Polymerase, 0.5 μM of forward and reverse primer and nuclease-free water. Cycling conditions included an initial denaturation at 95°C for 1 minute followed by 32 cycles of 95°C for 15 seconds, 59.5°C for 15 seconds and 72°C for 10 seconds and a final extension at 72°C for one minute in a thermocycler (Bio-Rad 382 T100 Thermal Cycler, USA). A non-template control (no DNA template) was included in each run and a 2% agarose gel was run to confirm the presence of a band of the correct size (382 bp).

Based on results from Experiment 1, a dose titration trial was performed (Experiment 2) using 10- fold dilutions of isolate Vh-14 such that the final doses were 4.0 × 10^8^, 3.0 × 10^7^, 3.0 × 10^6^ and 3.2 × 10^5^ CFU mL^-1^. Fish in Experiment 2 had a mean weight of 22.8 g ± 5.7 g (mean + SD) and a total length of 127.8 mm ± 11.6 mm. Fish were anaesthetised with benzocaine (60 mg L^-1^) prior to IM injection and were injected with 50 μL of Vh-14 adjusted for a final inoculum per fish between 10^4^-10^7^ CFU fish^-1^. The experimental design consisted of two replicate tanks for each challenge dose and one tank for the negative control (sham inoculation with PBS) and each tank contained 10 fish. After monitoring the fish for eight days, on Day 9, remaining fish were euthanised, and up to 4 fish per tank were randomly sampled to assess infection by culturing head-kidney swabs on TCBS plates followed by colony PCR as described for Experiment 1.

### 2.5 Statistical analysis

For infrared biotyping and the measurements from the IR Biotyper system, principal coordinate analysis (PCoA) was performed in R v4.2.1 [60] using cmdscale in stats 3.6.2 [61] and ggplot2 v3.5.1 [62] for visualisation. For experimental challenges, survival analysis was conducted using the Kaplan-Meier method and comparisons between survival curves made using Cox proportional hazard models followed by log-rank test. Moribund or dead fish were defined as experiencing the event of interest if they had clinical signs consistent with vibriosis and growth from a head-kidney swab on TCBS that was confirmed as *V. harveyi* by colony PCR. For Experiment 1, which tested different *V. harveyi* isolates at one dose (∼ 3.2 x 10^8^ CFU mL^-1^), isolate ID was fitted as a fixed effect. For the second experiment, which was the dose titration study of Vh-14, the Cox proportional hazard model was fitted with dose as a fixed effect and tank as a random clustering factor to incorporate tank variation into the model. All analysis was performed in in R v4.2.1 and survival analysis was conducted with packages survminer v0.5.0 [63], survival v3.6.4 [64], coxme v2.2.20 [65].

## 3. Results

### 3.1 *In silico* predictors of virulence

MALDI-ToF MS identified all newly obtained isolates sequenced in this study as *V. harveyi* (S4 Table) however average nucleotide identity (ANI) analysis showed that one isolate (Vh-24) had a 97.3% ANI to the *V. campbellii* reference strain (97.3% for BoB-53 vs 89.9% for SB1). The remaining isolates were confirmed as *V. harveyi* with > 97% ANI to SB1. Given the disparities in identification, the twenty-eight genomes sourced from the SRA were also checked with fastANI and three of the twenty-eight were re-classified as *V. campbellii* under fastANI: MWU1063 (SRR25169360), CUB2 (SRR2931635) and RT-6 (SRR5410471). Inclusion of these isolates for phylogenetic investigation resulted in a distinct outgroup from the other strains (S1 Fig) and recombination could not be calculated for these isolates due to the significant distances between SNPs when compared to the other isolates. Vh-24 was therefore excluded from phenotypic and *in vivo* study and all *V. campbellii* genomes were removed from phylogenetic and virulence gene investigation resulting in 37 genomes remaining, including the *V. harveyi* reference genome SB1. For short-read sequencing of the *V. harveyi* isolates collected from Vietnam, the mean number of bases per isolate was 5,863,706 (± 250,299) with a mean N50 of 378,834 bp and a high level of coverage (mean 93x; S4 Table). For long-read sequencing, the total number of bases sequenced was 7.23 Gb across 1,945,996 reads with an average genome size of 5,870,692 bp (± 154,712) across the isolates. The N50 read length was 6,850 bp with a median read length of 2,145 bp and a median Phred score of 19.34. The average GC content was 45% for both short and long-read sequencing methods. Vh-14 and Vh-15 had larger genomes than those of the other isolates from Vietnam for both short (average 6,277,849 ± 4302 bp for Vh-14 and Vh-15 versus 5,713,333 ± 32,459 bp for the rest, respectively) and long-read sequencing (average 6,095,344 ± 569 bp versus 5,778,011 ± 22,549 bp, respectively), due to the presence of additional plasmids (S4 Table).

Plasmids > 5000 bp in length were detected via long-read sequencing in three of the isolates: Vh- 14, Vh-15 and Vh-31 (S5 Table). In Vh-14 and Vh-15, eight and nine plasmids larger than 5,000 bp were assembled with the long-read data, respectively, and confirmed with Plassembler. Whilst four of the plasmids in Vh-14 were circular according to Plassember, only one small 5,097 bp sequence was circular in Vh-15. In addition, Plassembler identified two circular plasmids of 61,899 and 24,313 bp in Vh-31. In the absence of circularisation, a match between each plasmid and the plasmid database (PLSDB) [66, 67] with a mash distance of 0.1 or less was considered a key criterion for inclusion as a true plasmid [41]. Vh-14 had six hits to five distinct plasmids in PLSDB whilst Vh-15 had five hits to three plasmids (S5 Table). In Vh-14 the largest non-circular plasmid with a match in PLSDB was 105,412 bp in length and contained 131 coding sequences and two non-coding RNAs encoding for the *sok* antitoxin regulator, which is involved in plasmid maintenance and stability in conjunction with the *hok* gene [68].

All *V. harveyi* isolates from Vietnam sequenced in this study contained the same three antimicrobial resistance genes (S4 Table): *bla*(VHH-1), a type A beta-lactamase that facilitates hydrolysis of beta-lactams [69]; Tet(34), which is thought to disrupt tetracycline activity via activation of Mg^2+-^dependent purine nucleotide synthesis [70]; and Tet(35), an efflux pump involved in transporting tetracycline out of the cells [71]. The proportion of prophage content was variable across the isolates yet all isolates except Vh-33 contained at least 1 putative prophage region (S6 Table). Vh-14 and Vh-15 had the highest number of chromosomally dispersed prophage regions with three intact phage regions each.

Of the 49 protein-encoding virulence genes that were investigated, 38 were detected in all *V. harveyi* genomes analysed, indicating a large proportion of virulence genes are conserved (Fig 2A), while three atypical virulence genes originally identified in *V. cholerae* (*ctxA*, *tcpA*) and *V. parahaemolyticus* (*trh*) were not identified in any of the genomes. Four virulence genes were detected in one or two genomes only, including *vhml* (SRR11841529 and SRR17393361), *hylA* (SRR11841535), *vvh* (SRR11841535) and *ypkA* (*yopO*) (Vh-14 and Vh-15). There were five genes with more variable presence across the genomes including *zot* (present in 24 isolates), *ureB* and *ureG*, which encode for urease production (present in 32 isolates), and *virD4* (present in 22 isolates). The average number of virulence genes present in each genome was 41 (± 1) and all genomes in this study encoded a type III secretion system on Chromosome I (*vscN, vscF, vscB, vcrD* and *vcrH*). In addition to this chromosomal type III secretion system, Vh-14 and Vh-15 were the only two to encode *ypkA* – a serine/threonine kinase and effector protein within an additional type III secretion system (Lee et al., 2017). The 2,142 bp *ypkA* gene was encoded on the large 105,412 bp conjugative plasmid of Vh-14 and was also identified in a smaller fragmented plasmid in Vh-15 (Fig 3A). *YpkA* is an essential virulence gene of the *Yersinia* genus [72]. Several additional genes involved in the type III secretion system, *yopH, lcrG* and *lcrQ* and a type III chaperone were identified through manual screening against the *Yersinia* spp. reference genomes (Fig 3B). Surrounding the type III secretion system were numerous transposases and an integrase which can assist in the uptake and stabilisation of genes within bacterial plasmids [73].

**Fig 2.**
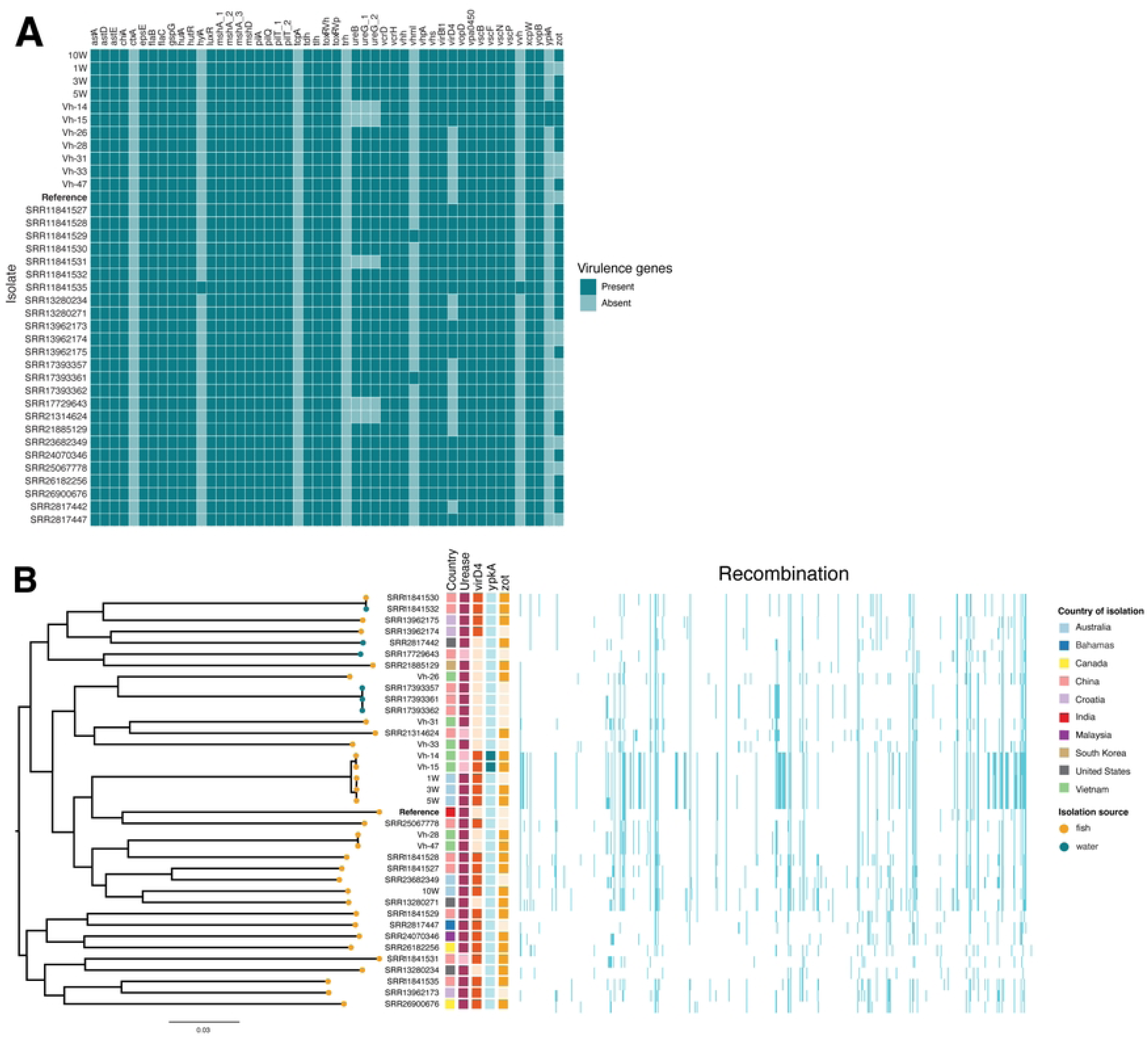
*In silico* virulence gene expression and single nucleotide polymorphism phylogeny. A) Presence-absence matrix for the occurrence of 49 protein-encoding virulence genes across 37 *Vibrio harveyi* genomes including the *V. harveyi* reference genome (SB1, NCBI: PRJNA972608; **bolded**). BLASTx was performed to determine whether a gene was present or absent within the genomes with a cut-off e-value of < 0.00001. B) Maximum-likelihood phylogenetic tree of 37 *V. harveyi* genomes. Tip colour indicates source of isolate (fish or environmental) and external blocks indicate country of origin, presence of *ureB/ureG (*Urease), *virD4, ypkA* and *zot* genes (indicated by darker shading) and homologous recombination, scale bar indicates average number of base substitutions per 1000 nucleotides. Figures generated using ggplot in R and Microreact (https://microreact.org/).

**Fig 3.**
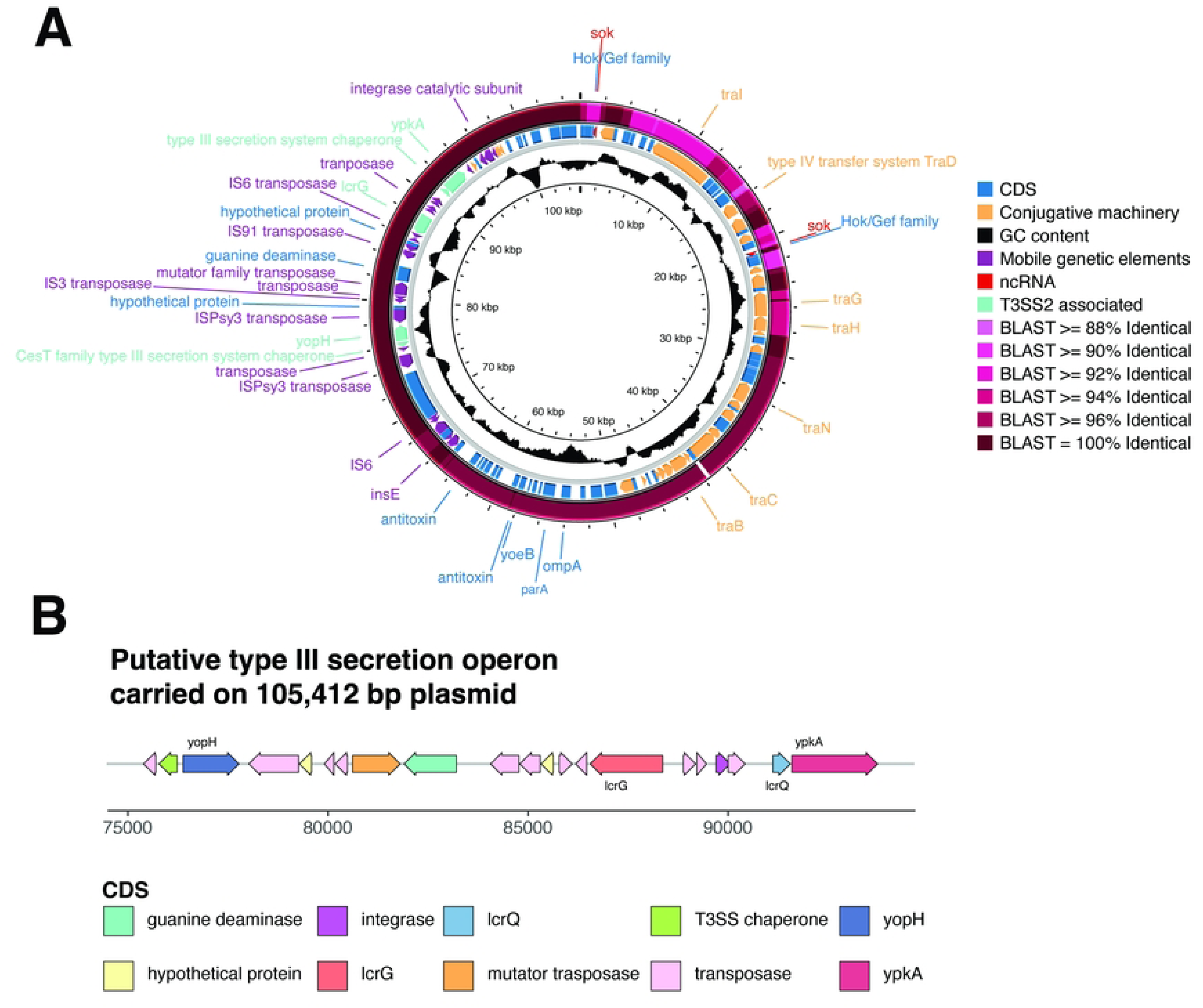
Plasmid identified in Vh-14 and Vh-15. A). Map of 105,412 bp plasmid identified in Vh-14. Major genes of interest are labelled, and colours correspond to the legend on the right of the circular map. An additional blastn was performed against Vh-15 and is the outermost ring on the map with identity to Vh-14 shaded in pink .Map was generated in Proksee with genes annotated via Bakta or manual BLASTx investigation (https://proksee.ca, [50]). B) Annotated map of genes of interest carried on 105,412 bp plasmid in Vh-14 and Vh-15 which contains components of a second type III secretion system. Genes annotated with Bakta or via manual BLASTx investigation of the plasmid against the *Yersinia* spp. reference genomes. Protein-coding genes of interest (*yopH, lcrG, lcrQ, ypkA*) are labelled and putative function of these genes and the surrounding ones are colour-coded. Figure generated in gggenes package in R.

Whilst isolates collected from the same disease investigations and hence typically the same site clustered together on the phylogeny, there was no other clear geographical pattern to phylogenetic relations (Fig 2B). Two of the isolates collected in Vietnam (Vh-14 and Vh-15) clustered together with three isolates sequenced from moribund barramundi in Australia (1W, 3W and 5W) and formed a sub-clade with the reference (SB1) genome and SRR25067778. When compared to the reference genome, Vietnamese isolates Vh-14, Vh-15 and the three Australia genomes (1W, 3W, 5W) had large levels of genomic recombination compared to the reference SB1.

### 3.2 *In vitro* virulence characteristics

For most phenotypic tests, including for caseinase, gelatinase, lipase, and phospholipase activity and swarming, there was an evident effect of temperature on observed activity, which was typically higher at 28°C or 34°C than at 22°C, (Fig 4A). Interestingly, biofilm production was the inverse with the highest levels of biofilm formation observed at 22°C for five of the eight *V. harveyi* isolates (S7 Table). Although there was overall increase in activity at higher temperatures, a large degree of individual variation was observed between isolates. This was most evident with haemolysis with only *P. aeruginosa* (the positive control) and Vh-31 producing complete β-haemolysis on sheep blood agar after 72 hours. For negative controls, there was no growth or activity with the exception of *E. coli* at 28°C on casein-specific media. At 28°C, small amounts of *E. coli* growth were observed below the agar following stabbing but without the formation of true clearing zones and hence not considered true caseinase activity. Urease activity was observed in all isolates and at all temperatures with the exception of Vh-14 and Vh-15 (S7 Table). These isolates were urease negative at all three temperatures and lacked urease genes: *ureB*, *ureG_1* and *ureG_2* found in the other isolates *in silico*.

**Fig 4.**
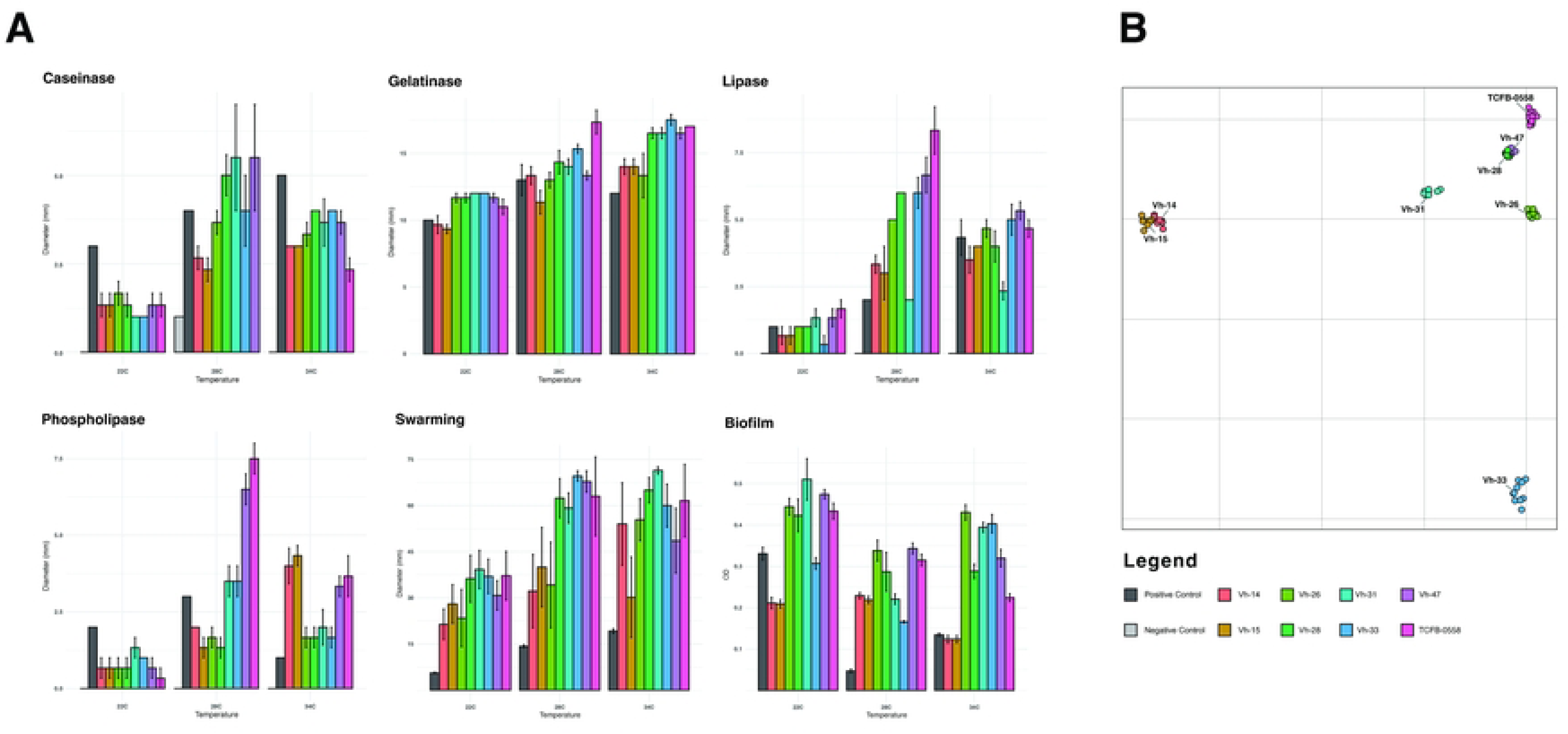
Phenotypic activity of *Vibrio harveyi* isolates. A) Bar plots depicting the variation in phenotypic activity of *Vibrio harveyi* isolates at three temperatures (22°C, 28°C and 34°C). Bar colours indicate different isolates on the x-axis; measurements of interest on the y-axis are: diameter in mm of clearing or opalescent zones for caseinase, gelatinase, lipase, phospholipase and swarming assays and OD_590nm_ for biofilm formation). Bar height is representative of the mean measurement of three replicates (five for swarming) with standard error bars. Where no negative control bar (light grey) can be seen, results were negative. B) Principal Coordinate Analysis plot of *V. harveyi* isolates investigated with the IR Biotyper system (Bruker Daltonics). Each isolate was typed in triplicate, by performing the run on three separate days (biological) with three replicates (technical) on each day.

Fourier Transform Infrared Spectroscopy was performed to assess isolates’ distinctive ‘fingerprints’. Similar to the findings of the genomic and other phenotypic analyses, Vh-14 and Vh-15 were highly distinctive (Fig 4B). Based on PCoA, they were highly similar to each other with an average distance of 6.30 to one another, below the IR Biotyper determined clustering cut-off of 6.93, but with an average Euclidean distance of 109.87 to other isolates (S8 Table). Vh-33 was also distinct from the other isolates with an average distance of 85.49. The remaining isolates clustered together in one cluster with an average distance of 35.98 from one another. Nested within this was a subcluster formed by Vh-28 and Vh-47 with an average distance of 3.38, below the IR biotyper clustering cut-off.

### 3.3 *In vivo* experimental challenges

In Experiment 1 (n = 2 replicates per isolate), there was significant (p < 0.001) and rapid mortality in juvenile barramundi injected with Vh-14 and Vh-15 with 100% cumulative mortality in less than 48 hours (Fig 5A) regardless of fish size. Significant cumulative mortality (p-value < 0.001) was also observed in fish infected with TCFB-0558 (positive control) with 70 ±10% standard error (mean ± SE), although mortality with this isolate was lower in large fish than small fish (p-value = 0.006, pairwise log-rank test). Several isolates caused low level mortality *in vivo,* e.g. a cumulative mortality of 9.5 ± 6.4% was observed for Vh-33, but this was not significantly different from the negative control (p = 0.24, pairwise log-rank test). Lower mortalities (one fish recorded for Vh-26 and Vh-47) were also not significantly different (p = 0.40) from the negative control group. No mortalities were recorded for Vh-28 or Vh-31 across the replicates. In Replicate 2, one fish from the negative control tanks was euthanised for animal welfare reasons after presenting skin lesions around the head consistent with inter-fish aggression and cannibalism in barramundi. The head-kidney of the euthanised control fish was swabbed onto TCBS and sheep blood TSA to confirm the absence of infection. No colonies grew from the head-kidney swab or from subsequent swabs from any of the control fish across all experiments.

**Fig 5.**
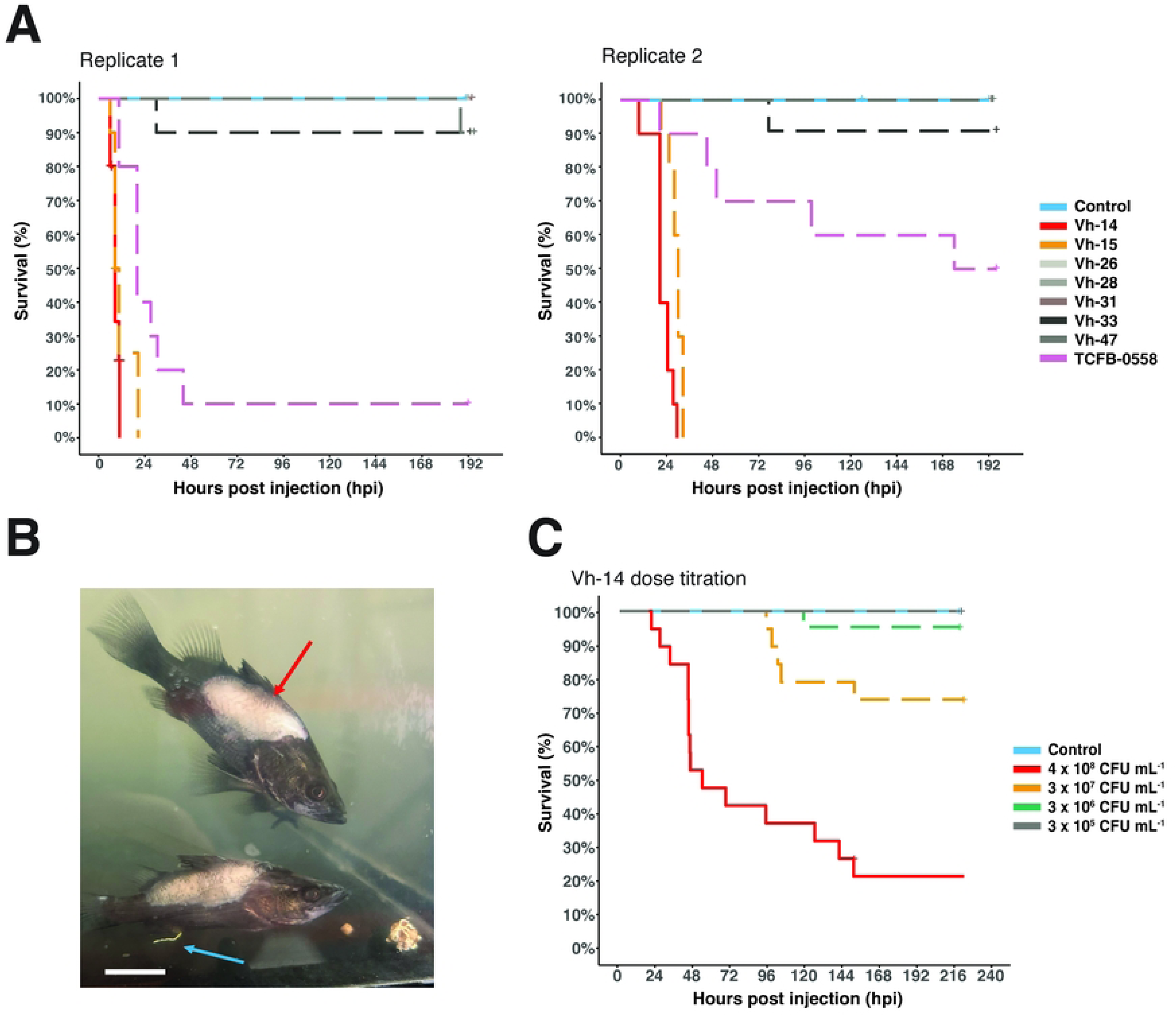
Kaplan-Meier survival curves for experimental challenge trials in barramundi. (A) Kaplan-Meier survival curves for intra-muscular (IM) challenge trials with eight *Vibrio harveyi* isolates in juvenile barramundi (*Lates calcarifer*). Control fish (n = 20) were IM injected with sterile PBS, whilst *V. harveyi* challenged fish (n = 10 /tank per replicate) were IM injected with ca. 3.2 × 10^8^ CFU mL^-1^. (B) Fish experiencing scale loss (red arrow) and gastroenteritis, visible as white casts (blue arrow), 3 days post-challenge in Experiment 1 (replicate 1), scale bar = 2 cm. (C) Kaplan-Meier survival curves for Experiment 2, with IM challenge doses of 10^5^ to 10^8^ CFU mL^-1^ of isolate Vh-14. Censored events such as survival or a lack of *V. harveyi* confirmed infection are indicated by a ‘+’.

Across all the isolates tested *in vivo*, clinical signs were observed as early as six hours post injection and continued until the conclusion of the observation period on Day 8. Clinical signs were highly varied between individuals and included scale loss, skin discolouration, lethargy, inappetence and erratic swimming. As infection progressed, increased severity of the above clinical signs and additional ones such as gastroenteritis (evidenced by white casts protruding from the vent, Fig 5B), skin and fin ulceration, and a loss of equilibrium were observed (Fig 5B). Ulceration and scale loss continued and resulted in the sloughing off of skin and muscle from 3 days post-injection. Despite the skin sloughing and exposure of the subcutaneous tissue, most fish continued to eat and swim normally and during the observation period were actively responding to stimuli. At the completion of the trial, many fish had experienced substantial wound-healing of their lesions. There was an apparent effect of fish length and weight on mortality kinetics with smaller fish experiencing more rapid mortality compared to larger fish. Across the two replicates, the nature and onset of clinical signs was consistent across fish sizes as was mortality for each isolate.

In Experiment 1, head-kidney swabs from 53 of the 56 (95%) moribund and dead fish yielded growth on TCBS plates, with characteristic yellow, smooth and round colonies that were all confirmed as *V. harveyi* by conventional PCR. The three fish (two injected with Vh-14 and one injected with Vh-15) that did not develop positive head-kidney swabs died within the first 12 hours following challenge, displaying behavioural and clinical signs consistent with vibriosis (keeling on their side, unresponsive to stimuli, darker coloration of the skin). Of the head-kidney swabs collected at the completion of the trial, 93% of fish (41/44) presented growth on TCBS plates, with 87% (36/41) confirmed as *V. harveyi* by conventional PCR.

Experiment 2 investigated the dose kinetics of Vh-14 based on the rapid and high cumulative mortality observed for Vh-14 in Experiment 1. In comparison to our prior challenge model study with TCFB-0558 [21], Vh-14 caused mortality at lower doses with significantly lower survival for Doses 1, 2 and 3 (p-value < 0.001) in comparison to the negative control treatment (Fig 5C). For Dose 1 (4.0 × 10^8^ CFU mL^-1^), total cumulative mortality was 79 ± 9.4% across the trial and mortality occurred later than in the smaller fish used in Experiment 1. Mortalities were lower (26 ± 10% and 5 ± 4.6%, respectively) and occurred later (4- and 5-days post injection) for Dose 2 (3.0 × 10^7^ CFU mL^-1^) and Dose 3 (3.0 × 10^6^ CFU mL^-1^) compared to Dose 1. Clinical signs were consistent with those observed in Experiment 1 and additional gross internal pathology was observed at the conclusion of the trial including splenomegaly and discoloration of the liver in fish receiving Dose 1 or 2. Fish injected with Dose 4 (3.2 × 10^5^ CFU mL^-1^) experienced some lethargy following injection but exhibited no other clinical signs and were responsive and feeding within 24 hours with no detectable *V. harveyi* in the head-kidney or skin at the completion of the trial. All moribund fish and dead fish recorded in Experiment 2 were positive for *V. harveyi* by culture and conventional PCR, with the exception of one fish that died overnight and had started decomposing, resulting in release of gastrointestinal bacteria. At the end of the trial, all fish sampled in Dose 1 and Dose 2 had growth on TCBS plates that was confirmed by PCR as *V. harveyi*, except for one head-kidney swab taken from a fish exposed to Dose 2. For Doses 3 and 4, 37.5% (6/16) of swabs taken from fish yielded growth on TCBS plates with characteristic colonies, but only one from Dose 3 was confirmed as *V. harveyi*.

## 4. Discussion

In the face of climate change and rising sea-surface temperatures, an increase in the prevalence of *Vibrio* spp. worldwide is already occurring [74, 75]. Increased attention to the threat of *Vibrio* spp. that are relevant to public health, such as *V. vulnificus* [76], non-O1 *V. cholerae* [3] and *V. parahaemolyticus* [77] is occurring as human case numbers continue to rise. By contrast, the potential impacts of *Vibrio* spp. on sustainable aquaculture and food security have received limited attention, despite the recent recognition of this genus as a dominant bacterial agent associated with disease and mortality in finfish and crustacean aquaculture globally [10, 13, 46]. Although some virulence characteristics of *V. harveyi* have been described [78], it is still unclear how or why certain strains cause severe disease while others remain avirulent unless the host experiences considerable stress [21, 79]. Our study sought to integrate genomic analyses, conventional phenotypic assays and *in vivo* trials in barramundi fingerlings to expand understanding of *V. harveyi* pathogenesis.

Across all isolates, activity deemed indicative of virulence was increased at higher temperatures with the exception of urease activity and biofilm formation. Whether this effect is mediated simply by preferred growth conditions or virulence gene expression remains unclear. Evidence of increased growth rates within the first 48 hours at higher temperatures was observed for *V. harveyi* cultured in seawater at 25°C and 30°C compared to 20°C [80] and at 30°C compared to 26°C [81], however, this could also be associated with decrease in culturability in seawater after two or more weeks. Transcriptomic analysis is suggestive of higher virulence gene expression for *V. harveyi* at 30°C with increased expression of chemotaxis and several type III and IV secretion system proteins compared to 26°C [81]. This change in gene expression has also been observed *in vitro* for *V. vulnificus* at higher temperatures (28°C vs 20°C) resulting in the activation of metabolic components such as chemotaxis and proteases to assist in host colonisation within European eels (*Anguilla anguilla*) [82]. Increases in temperature could therefore act as a stressor and in conjunction with other stressors such as high salinity or pH contribute to the increased secretion of virulence components and host adaptation as previously shown for the food-borne *Enterococcus* spp. [83] and the fish pathogen *Aeromonas hydrophila* [84]. Some *in vivo* effect of temperature on virulence has also been observed with an increase in mortality of red tilapia (*Oreochromis* sp.) challenged with *Streptococcus agalactiae* [85]. Similar effects have been observed in Pacific oysters (*Crassostrea gigas*) with an increase in mortality and the presence of numerous *Vibrio* spp. in the microbiome during simulated marine heatwaves [86] and during a summer mortality outbreak [87]. The effect of thermal stressors on increasing virulence are supported by predictive modelling showing that an increase of just 1°C environmentally could lead to a 3.5% increase in mortality of aquatic organisms [88]. Despite the expression of virulence characteristics *in vitro,* the majority of isolates did not cause mortality in our challenge study, although they may have contributed to the disease syndromes in farmed marine fish in Vietnam observed during the isolation process. In addition to exerting pressure on the pathogen, stressors also play a major role in host response and susceptibility to *V. harveyi.* Typically co-stressors such as skin lesions [89] or thermal stressors [21] are necessary for disease to occur in otherwise healthy animals. Without testing the effect of these stressors alone or in combination, experimental infection models cannot always replicate the mortality kinetics observed on farm. Given that the Quang Ninh province in the North of Vietnam is a key tourism area and experiences heavy anthropogenic impacts within the surrounding regions (also reporting increased eutrophication [90] and microbial contamination [91]), finfish in this region most likely experience significant environmental stress facilitating infection and disease progression in the field. Interestingly, fish challenged experimentally with Vh-14 and Vh-15 did experience 100% mortality at a high dose (ca. 3.5 × 10^8^ CFU mL^-1^) without any additional stressors. Fish challenged with Vh-14 also exhibited dose-dependent mortality kinetics, not seen in our previous work with TCFB-0558 [21], suggesting a potentially highly virulent isolate capable of causing disease without the need for co-participating factors when injected in a sufficient dose.

Isolates Vh-14 and Vh-15 were distinctive *in vivo* as well as *in vitro* and *in silico*, in that they were urease negative and lacked the genes for *ureB* and *ureG1*/*2*. Urease activity is typical of the *V. harveyi* clade with reports that upwards of 50% of isolates are urease positive [92–94]. In the highly specialised fish pathogen *Photobacterium damselae* subsp*. piscicida,* the loss of urease catalysation resulted in an increase in host adaption and virulence from the generalist *P. damselae* subsp. *damselae* [95]. This shedding of non-essential genes including urease has also been observed in the evolution of *Y. pestis* from *Y. pseudotuberculosis,* facilitating greater transmission via the flea vector [96].

Whilst all *V. harveyi* genomes analysed carried a type III secretion system 1 (T3SS1: *vopD, vscB/F/N/P, vcrD/H)* on chromosome I [97], hypervirulent isolates Vh-14 and Vh-15 maintained a secondary plasmid-associated type III secretion system. Highly similar to the type III secretion system first identified in *Yersinia* spp. [98], this type III secretion system is also the major virulence mechanism of the economically important aquaculture pathogen *Aeromonas salmonicida* [99, 100]. A homologue of this additional type III secretion system has been identified in *V. cholerae* and *V. parahaemolyticus* [101] as type III secretion system 2 (T3SS2), although a universal nomenclature for the genes involved in the T3SS2 in *Vibrio* spp. is yet to be developed [102].

Upon first contact with host cells [103], the six effector proteins of the T3SS2 are responsible for the prevention of phagocytosis by the host [104]. Each of the effector proteins performs this is in a distinct manner with YpkA responsible for actin-filament disruption, impairing the cytoskeletal function [105] whilst YopH is antiphagocytic via inactivating neutrophils and by dephosphorylating focal adhesion complexes [106]. The other two protein-encoding genes *lcrQ* and *lcrG* identified in Vh-14 and Vh-15 are negative regulators of the effector proteins and needle apparatus contributing to cytoplasm stability in the bacteria [107].

Whilst plasmids have previously been detected in the genomes of some *V. harveyi* isolates [48, 49, 108, 109], there is currently no identified virulence plasmid of *V. harveyi.* Using long-read sequencing, our study found a conjugative plasmid in two isolates carrying components of a secondary type III secretion system suggesting the emergence of a hypervirulent plasmid-mediated patho-variant of *V. harveyi* with implications for marine finfish aquaculture. The assembly of these large plasmids carrying multiple repeated elements (eg. transposons, insertion sequences) is generally not possible with short-read sequencing alone [110], highlighting the critical role of long-read sequencing technology. In public health investigations, long-read sequencing is becoming an increasingly used tool for the monitoring of hypervirulent plasmids in multi-host pathogens including *Klebsiella pneumoniae* [111, 112] and group B *Streptococcus* (*Streptococcus agalactiae*), which affects aquatic and terrestrial species [113]. In aquatic systems, a key example of plasmid-mediated virulence has been documented for acute hepatopancreatic necrosis disease (APHND) in whiteleg shrimp (*Penaeus vannamei*) caused by the hypervirulent pVA1-type *V. parahaemolyticus* plasmid, which subsequently spread to other *Vibrio* spp. including *V. campbellii* [114]. Marine environments are a hotspot for horizontal gene transfer, with higher gene transfer rates observed for bacteria co-existing with archaea in high temperature environments [115] and within ocean environments [116]. As such, the increased use of long-read sequencing for *V. harveyi* and other aquatic bacteria of interest in aquaculture will assist with in the characterisation of previously unknown virulence components, improving our understanding of virulence evolution in key marine pathogens [117].

In conclusion, in this study we identified two potentially hypervirulent plasmid-mediated patho-variants of *V. harveyi* (Vh-14 and Vh-15) that encoded components of an additional type III secretion system on a conjugative plasmid. Further investigation into the role of this plasmid, its prevalence in the global aquaculture industry, and the likelihood of conjugation and gene transfer to other *V. harveyi* strains under a range of environmental conditions is crucial given the potential drastic effect on fish mortality in farmed settings were such patho-variants to become widespread.

## Animal Ethics Statement

All animal procedures were approved by The University of Sydney Animal Ethics Committee (AEC) under AEC project number 2023/2397 according to the Australian Code of Practice for the Care and Use of Animals for Scientific Purposes (8th edition, 2013).

## Author Contributions

RS: Conceptualisation, Investigation, Methodology, Data Curation, Formal Analysis, Visualisation, Writing – Original Draft, Writing – Review and Editing.

JAB: Conceptualisation, Investigation, Methodology, Funding Acquisition, Resources, Supervision, Writing – Review and Editing.

RNZ: Conceptualisation, Investigation, Methodology, Resources, Supervision, Writing – Review and Editing.

CV: Investigation, Methodology, Funding Acquisition, Writing – Review and Editing.

AE: Investigation, Methodology, Visualisation, Writing – Review and Editing.

SB: Investigation, Methodology, Data Curation, Writing – Review and Editing.

DLF: Investigation, Data Curation, Software, Writing – Review and Editing.

HA: Investigation, Data Curation, Software, Writing – Review and Editing.

GRO: Investigation, Data Curation, Software, Writing – Review and Editing.

VB: Funding Acquisition, Resources, Supervision, Writing – Review and Editing.

MS: Software, Methodology, Writing – Review and Editing.

MW: Supervision, Resources, Writing – Review and Editing.

PN: Investigation, Funding Acquisition, Writing – Review and Editing.

FS: Conceptualisation, Investigation, Methodology, Data Curation, Funding Acquisition, Software, Resources, Project Administration, Supervision, Writing – Review and Editing

## Data availability

Raw reads for both the Illumina short-read (paired) sequencing and Oxford nanopore long-read (single) sequencing have been deposited into the NCBI Sequence Read Archive under the Bioproject PRJNA1210070.

## Acknowledgements

The authors acknowledge the technical assistance provided by Stuart Glover, Jade Sutton, Cara Jeffrey Anna Waldron and Slavicka Patten. We acknowledge the efforts of Aakriti Regmi for assistance with sample preparation and experimental trial data entry. We thank Andrew Eliot at the Elizabeth Macarthur Agricultural Institute for providing access to incubators and Elise Beggs for her assistance in the lab. We acknowledge the Sydney Informatics Hub at the University of Sydney for providing access to high-performance computing and technical assistance, with special thanks to Georgie Samaha, Fred Jaya and Nandan Deshpande for their assistance and pipeline development.

## Supporting information

**S1 Appendix**. **Supplemental methods for Section 2.3.**

**S1 Fig. Phylogeny including mis-identified *V. campbellii*.** SNP-based phylogeny for the 12 isolates sequenced with short-read sequencing in this study (labelled Vh- or -W), the twenty-eight sequences downloaded from the SRA labelled with their unique SRR number and the *Vibrio harveyi* reference genome (SB1, NCBI: PRJNA972608). A distinct outgroup of four isolates (shaded in blue) formed from the other isolates. Vh-24 had an 97.3% ANI to the *Vibrio campbellii* reference genome (BoB-53, NCBI: PRJNA429202).

**S1 Table. 49 protein-encoding virulence genes identified in *V. harveyi* or other species of relevance.**

**S2 Table. Geographical and host origin of publicly available Vibrio harveyi genomes included for phylogenetic analyses.**

**S3 Table. Doses of V. harveyi determined using colony forming unit (CFU) counts performed retrospectively.**

**S4 Table. Summary of short and long-read sequencing for the isolates sequenced in the study.** Additional identification to a species level using MaLDi-TOF Matrix-assisted laser-desorption ionisation time-of-flight mass spectrometry (MALDI-ToF MS) and average nucleotide identity (ANI, as determined with FastANI) is included.

**S5 Table. Plassembler results for Vibrio spp. isolates containing plasmids detected during long-read sequencing with coding sequences and non-coding RNA annotated using Bakta.**

**S6 Table. Prophage regions predicted using PHASTEST in Vibrio harveyi isolates sequenced in this study.**

**S7 Table. Phenotypic characteristics of Vibrio harveyi at three different temperatures: 22C, 28°C (optimal growth temperature) and 34°C.** For Biofilm, mean OD + SE at 600nm is reported. For urease, a positive or negative result is reported and for haemolysis, a positive or negative result is reported and the mean + SE in mm is reported for those exhibiting haemolysis. All other tests are reported as the mean + SE in mm for all replicates for each isolate under each condition.

**S8 Table. Distance matrix generated from IR Biotyper software for Fourier Transform Infrared Spectroscopy for the eight V. harveyi isolates phenotypically investigated in this study.**

